# Local brain oscillations and inter-regional connectivity differentially serve sensory and expectation effects on pain

**DOI:** 10.1101/2022.08.10.503459

**Authors:** Felix S. Bott, Moritz M. Nickel, Vanessa D. Hohn, Elisabeth S. May, Cristina Gil Ávila, Laura Tiemann, Joachim Gross, Markus Ploner

## Abstract

Pain emerges from the integration of sensory information about threats and contextual information such as an individual’s expectations. However, how sensory and contextual effects on pain are served by the brain is not fully understood so far. To address this question, we applied brief painful stimuli to 40 healthy human participants and independently varied stimulus intensity and expectations. Concurrently, we recorded electroencephalography. We assessed local oscillatory brain activity and inter-regional functional connectivity in a network of six brain regions playing key roles in the processing of pain. We found that sensory information predominantly influenced local brain oscillations. In contrast, expectations exclusively influenced inter-regional connectivity. Specifically, expectations altered connectivity at alpha (8-12 Hz) frequencies from prefrontal to somatosensory cortex. Moreover, discrepancies between sensory information and expectations, i.e., prediction errors, influenced connectivity at gamma (60-100 Hz) frequencies. These findings reveal how fundamentally different brain mechanisms serve sensory and contextual effects on pain.

**Teaser:** Sensory and expectation effects on pain are implemented by fundamentally different brain mechanisms.

## Introduction

Pain serves to protect the body. To this end, the brain translates sensory information about potential threat into an unpleasant experience and protective behavioral responses. However, this translation is not only shaped by sensory but also by contextual information, such as an individual’s expectations (*1–3*). Expectations can yield powerful and clinically relevant changes of the pain experience, for example through placebo and nocebo effects (*4–7*). Moreover, contextual and expectation effects are particularly relevant for pathological aberrations of the pain experience in chronic pain disorders (*8–10*).

In the brain, pain is associated with complex patterns of neural activity in somatosensory, insular, cingulate, and prefrontal cortices as well as subcortical brain areas (*11, 12*). Neurophysiological studies using electroencephalography (EEG), magnetoencephalography (MEG), and intracranial recordings have shown that this brain network yields complex temporal-spectral patterns of neural responses including evoked potentials and oscillatory responses at alpha (8-12 Hz), beta (13-30 Hz), and gamma (30-100 Hz) frequencies (*13*). In addition, it is increasingly recognized that not only local brain activity, but also the communication between brain regions, i.e., interregional brain connectivity, critically shapes the experience of pain (*14–20*).

Recent studies have started to unravel how these complex spatial-temporal-spectral patterns of brain activity serve sensory and contextual effects on pain. Functional magnetic resonance imaging (fMRI) studies have revealed that these effects are served by different spatial patterns of brain activity. For instance, patterns of brain activity termed the neurological pain signature (NPS) and the stimulus intensity independent pain signature (SIIPS) are particularly sensitive to sensory and contextual effects on pain, respectively (*21, 22*). EEG studies have indicated that the temporal-spectral patterns of sensory and contextual effects on pain also differ (*23–25*). Specifically, evoked potentials and oscillatory responses to noxious stimuli are more sensitive to sensory information than to expectations (*23–25*). In contrast, the temporal-spectral pattern of expectation effects on pain has remained largely unclear so far. Mechanistic considerations suggest that contextual effects on pain such as expectations might be particularly shaped by inter-regional top-down connectivity between supra-modal and sensory brain regions at alpha and beta frequencies (*13*). Moreover, predictive coding frameworks of brain function (*26, 27*) propose that discrepancies between sensory and expectation effects, i.e., prediction errors (PE), are mediated by brain oscillations and connectivity at gamma frequencies (*28–30*). However, direct evidence for these hypotheses on how sensory and expectation effects on pain are implemented by local brain activity and inter-regional connectivity is lacking so far.

To better understand and directly compare how local brain oscillations and interregional connectivity serve sensory and contextual effects on pain, we re-analyzed data from an EEG experiment in which brief painful stimuli were applied to healthy human participants (*23*). Therein, sensory and contextual information was modulated by varying stimulus intensity and expectations about upcoming stimulus intensity, respectively. We here assessed and compared how local oscillatory brain activity and inter-regional connectivity in a core network of six brain regions associated with the processing of pain serves the effects of stimulus intensity and expectations on pain.

## Results

To investigate how the brain serves sensory and contextual influences on pain, we employed a probabilistic cueing paradigm. We applied brief painful heat stimuli to the left hand and independently modulated stimulus intensity and expectations in a 2 × 2 factorial design. To modulate stimulus intensity, we applied painful stimuli of two different levels (high intensity [hi] and low intensity [li]). To modulate expectations, the painful stimuli were preceded by one out of two visual cues, probabilistically indicating the intensity of the subsequent stimulus. The high expectation (HE) cue was followed by a hi stimulus in 75% of the trials and by a li stimulus in 25% of the trials. Conversely, the low expectation (LE) cue was followed by a hi stimulus in 25% of the trials and by a li stimulus in 75% of the trials. The experiment thus comprised four trial types (Figure 1A): high intensity, high expectation (hiHE); high intensity, low expectation (hiLE); low intensity, high expectation (liHE); low intensity, low expectation, (liLE). In each trial, after the painful stimulus, the participants were asked to provide a rating of the perceived pain intensity on a numerical rating scale ranging from 0 (no pain) to 100 (maximum tolerable pain). Figure 1b shows the sequence of a single trial. The experiment included 160 trials per participant.

**Figure 1.**
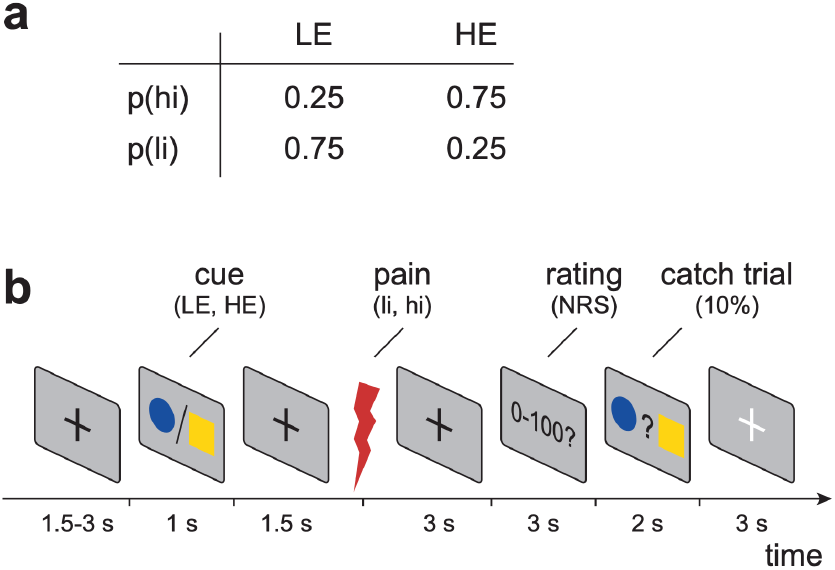
Experimental design. (a) Probabilities of different pain stimulus intensities (low intensity, li; high intensity, hi) for different levels of expectation (low expectation, LE; high expectation, HE). (b) In each trial, a cue was presented that probabilistically predicted the intensity of a subsequent painful stimulus. Three seconds after the stimulus, a verbal pain rating was obtained from the participants. More details on the experimental design can be found in [Nickel et al. 2022].

We analyzed oscillatory brain activity and functional connectivity in a network of six brain regions (Figure 2) known to play key roles in the cerebral processing of pain (*31*). The brain regions were the contralateral primary somatosensory cortex (S1), the contra- and ipsilateral parietal operculum (cPO, iPO; including the secondary somatosensory cortex and parts of the insular cortex), the anterior cingulate cortex (ACC), and the contra- and ipsilateral prefrontal cortex (cPFC, iPFC). Some of these brain regions are particularly associated with processing of sensory information (S1, cPO, iPO) whereas others are more associated with supramodal cognitive and emotional processes (ACC, cPFC, iPFC) (*11, 12*). Coordinates for these six regions of interest (ROIs) were taken from human intracranial recordings which represent the gold standard for electrophysiological brain responses to pain stimuli (*31*). To assess oscillatory brain activity, we calculated frequency-specific power in source space. To assess functional connectivity between brain regions, we calculated the debiased weighted phase lag index (dwPLI) (*32*). Both local activity and inter-regional connectivity were assessed in the alpha (8-12 Hz), beta (14-30 Hz), and gamma (60-100 Hz) frequency bands. These frequency bands are known to exhibit changes in oscillatory power in response to brief painful stimuli (*33–36*) and play key roles in interregional communication in the brain (*37*). In addition, to assess the dominant direction of information flow in selected connections and frequency bands, we computed an asymmetry index based on the partial directed coherence measure (*38*) of directed functional connectivity.

**Figure 2.**
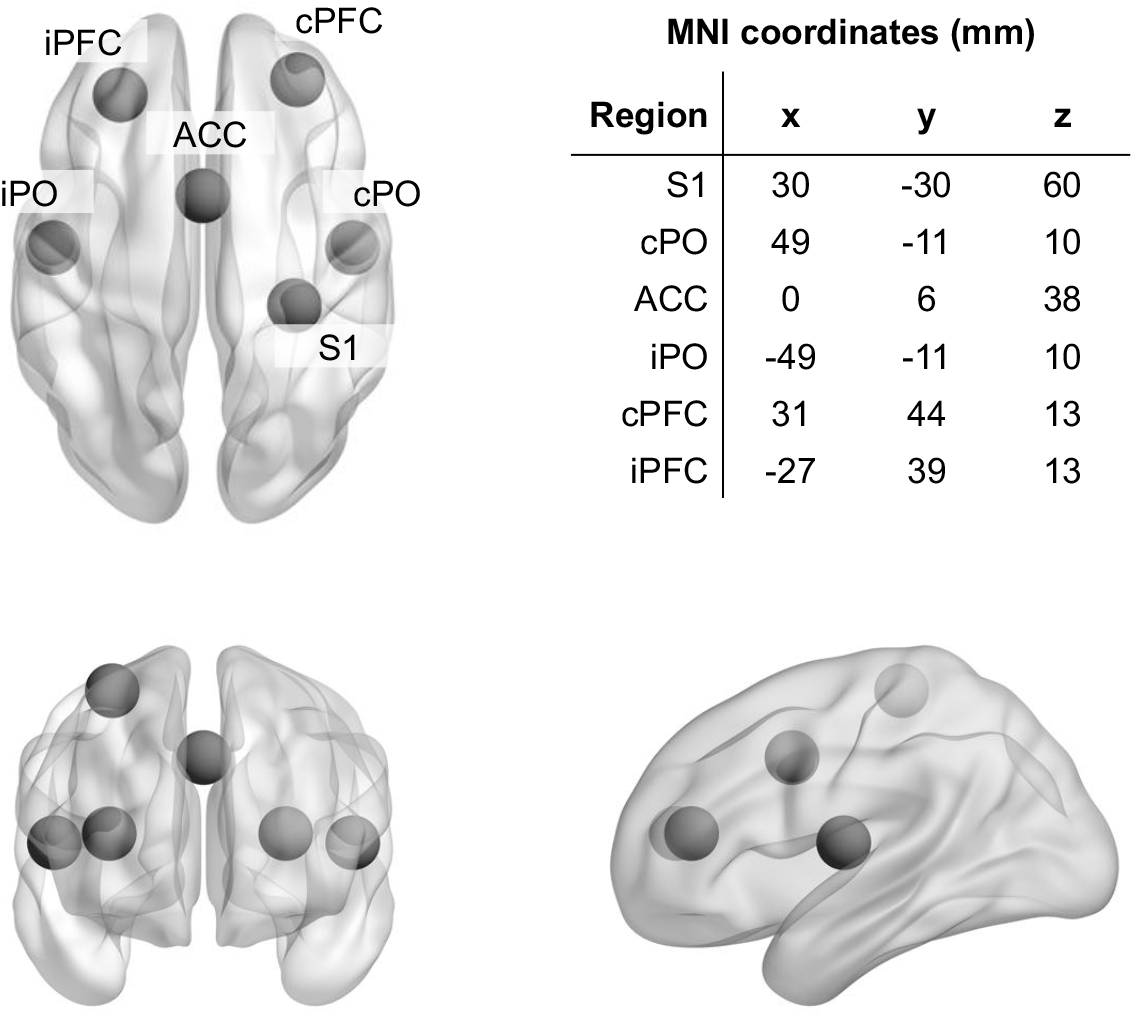
Regions of interest and corresponding MNI coordinates. Axial, coronal, and sagittal view of the brain and the six regions of interest. S1: contralateral primary somatosensory cortex; cPO, iPO: contra- and ipsilateral parietal operculum; ACC: anterior cingulate cortex; cPFC, iPFC: contra- and ipsilateral prefrontal cortex.

To relate brain activity and connectivity to sensory and expectation effects on pain, we defined different patterns describing the relation between response variables and experimental manipulations (*39, 40*). In particular, these patterns characterize how neural phenomena and pain ratings are linked to intensity, expectations, or discrepancies thereof (prediction errors, PEs) across the four trial types (Figure 3). To formally link the data to these patterns, we performed repeated measures analyses of variance (rmANOVAs) (*41*) with the independent variables intensity and expectation. In these rmANOVAs, features signaling stimulus intensity and expectations would manifest as main effects, whereas features signaling PEs would manifest as interactions. To allow for the interpretation of negative findings, we specifically performed Bayesian rmANOVAs (*41*).

**Figure 3.**
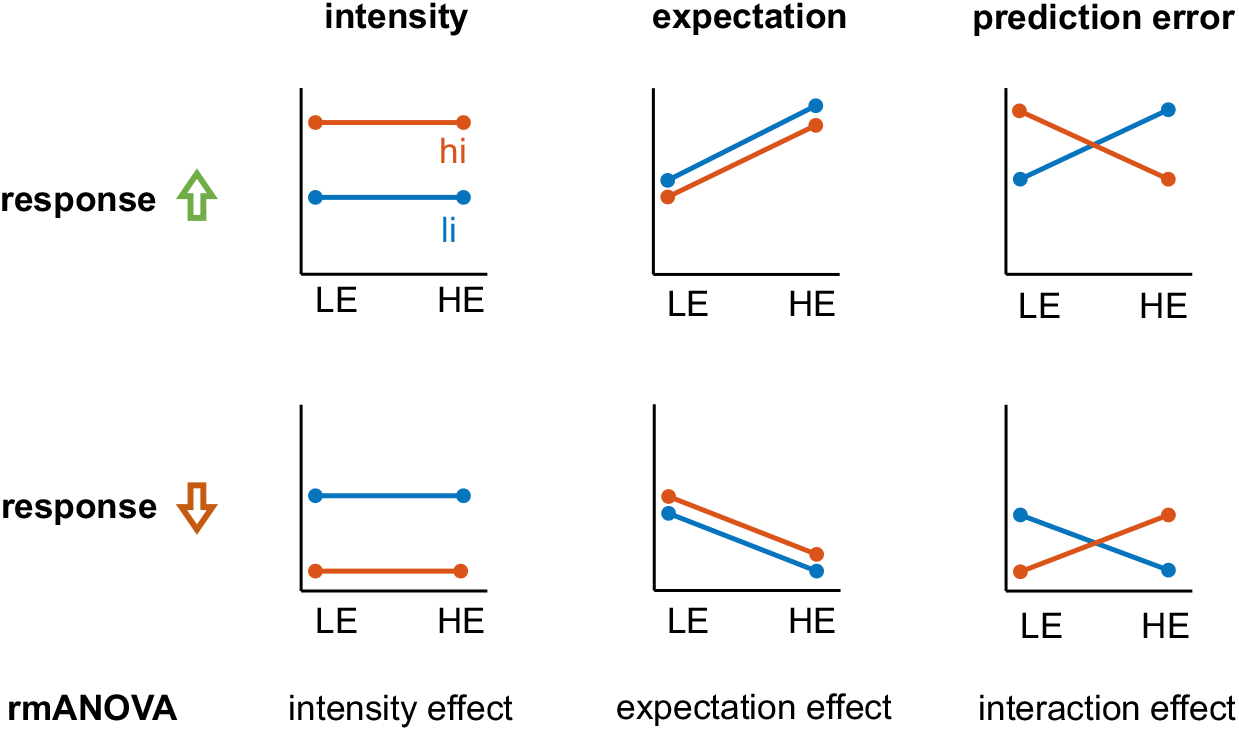
Possible response patterns indicating effects of stimulus intensity, expectations, and PEs. Effects of stimulus intensity (low intensity, li; high intensity, hi), expectations (low expectation, LE; high expectation, HE), and prediction errors were tested by means of rmANOVAs. An experimental modulation can lead to either a relative increase (first row) or relative decrease (second row) of oscillatory activity or connectivity.

### The effects of stimulus intensity and expectations on pain intensity ratings

Figure 4 shows pain intensity ratings for the four trial types. Analyses of pain ratings provided decisive evidence for main effects of intensity (BF = 1.1 *10^21^) and expectation (BF = 5.5*10^2^) on pain ratings. Specifically, as expected, hi stimuli yielded higher pain ratings than li stimuli, and HE cues yielded higher pain ratings than LE cues. Moreover, there was moderate evidence against an interaction effect of intensity and expectation (BF = 0.27). Thus, the results confirmed that stimulus intensity and expectations shaped pain ratings.

**Figure 4.**
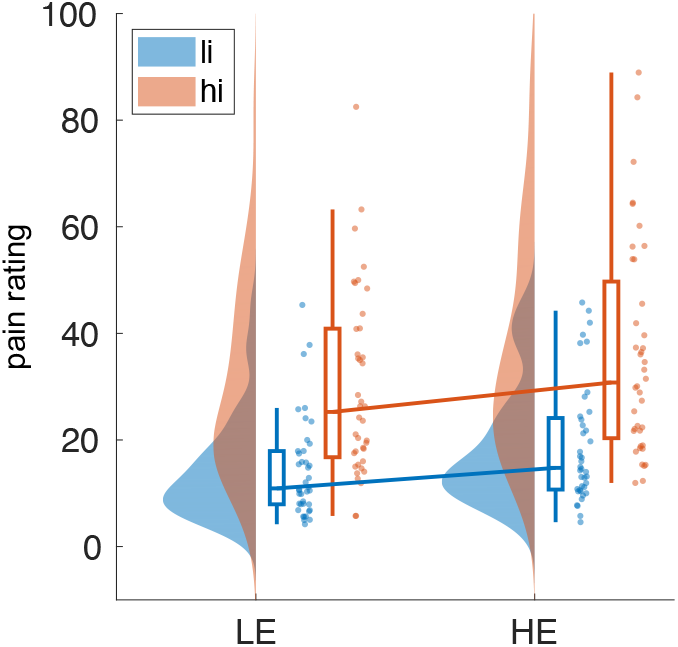
Effects of stimulus intensity, expectations, and prediction errors on pain ratings. Rain cloud plot (*42*) of pain ratings for two levels of stimulus intensity (low intensity, li; high intensity, hi) and expectation (low expectation, LE; high expectation, HE). A Bayesian rmANOVA yielded decisive evidence for main effects of stimulus intensity and expectation (BF = 1.1*10^21^ and BF = 5.5*10^2^, respectively). Moreover, there was moderate evidence against an interaction (BF = 0.27).

### The effects of stimulus intensity and expectations on local oscillatory brain activity

We first assessed how brief noxious stimuli influenced local oscillatory brain activity in the six ROIs. Time-frequency representations (TFRs, **Fehler! Verweisquelle konnte nicht gefunden werden.**) indicated that noxious stimuli suppressed alpha and beta activity in all ROIs and increased gamma activity predominantly in S1. In addition, noxious stimuli yielded increases of activity at frequencies below 8 Hz which reflect evoked potentials analyzed previously (*23*).

Next, we assessed how stimulus intensity and expectations influence local brain activity in our core network associated with pain processing. We therefore determined the power of brain activity in the six ROIs at alpha, beta, and gamma frequencies averaged across the 1 s post-stimulus interval. The results of Bayesian rmANOVAs with the factors intensity and expectation are shown in Figure 6. We found that stimulus intensity modulated local brain activity at all frequency bands and in all ROIs. Strongest stimulus intensity effects were observed at alpha frequencies where we found moderate to decisive evidence for effects on oscillatory brain activity for all ROIs. In all ROIs, stronger stimuli yielded stronger suppressions of alpha activity. Weaker effects were observed at beta frequencies where we found moderate evidence for an intensity effect on brain activity in S1, iPO, and cPFC. In these ROIs, stronger stimuli yielded stronger suppressions of beta activity. In the gamma frequency band, we observed moderate evidence for an intensity effect on S1 brain activity with stronger stimuli inducing higher amplitudes of gamma activity.

**Figure 5.**
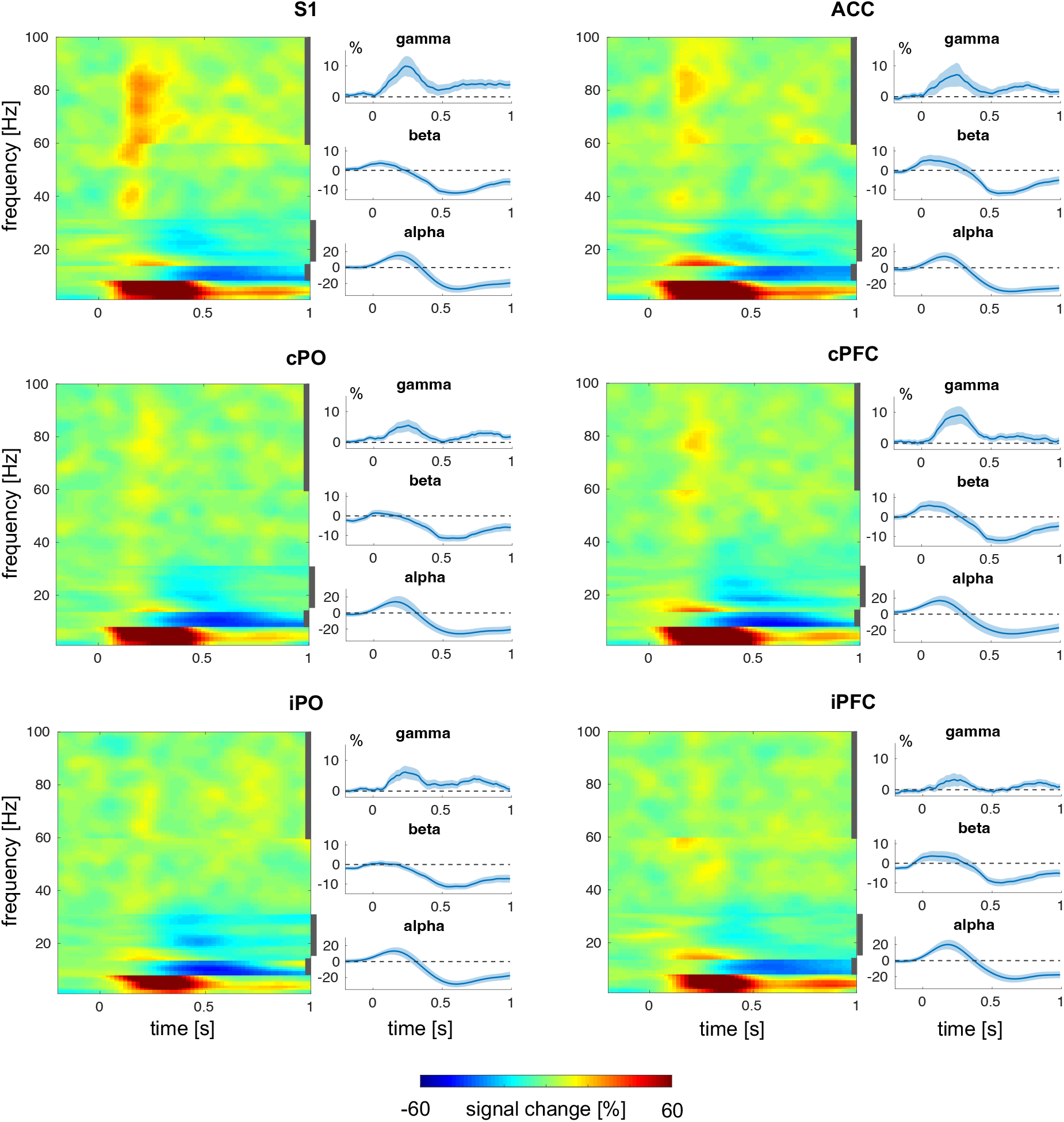
Time-frequency representations of local oscillatory brain activity in the six ROIs. The first and third columns show concatenated band specific TFRs for all six ROIs. The second and fourth columns show time-courses of brain activity in the alpha, beta and gamma band. Vertical, dark-gray bars in the TFR plots indicate the frequency intervals based on which the time courses of brain activity were computed.

**Figure 6.**
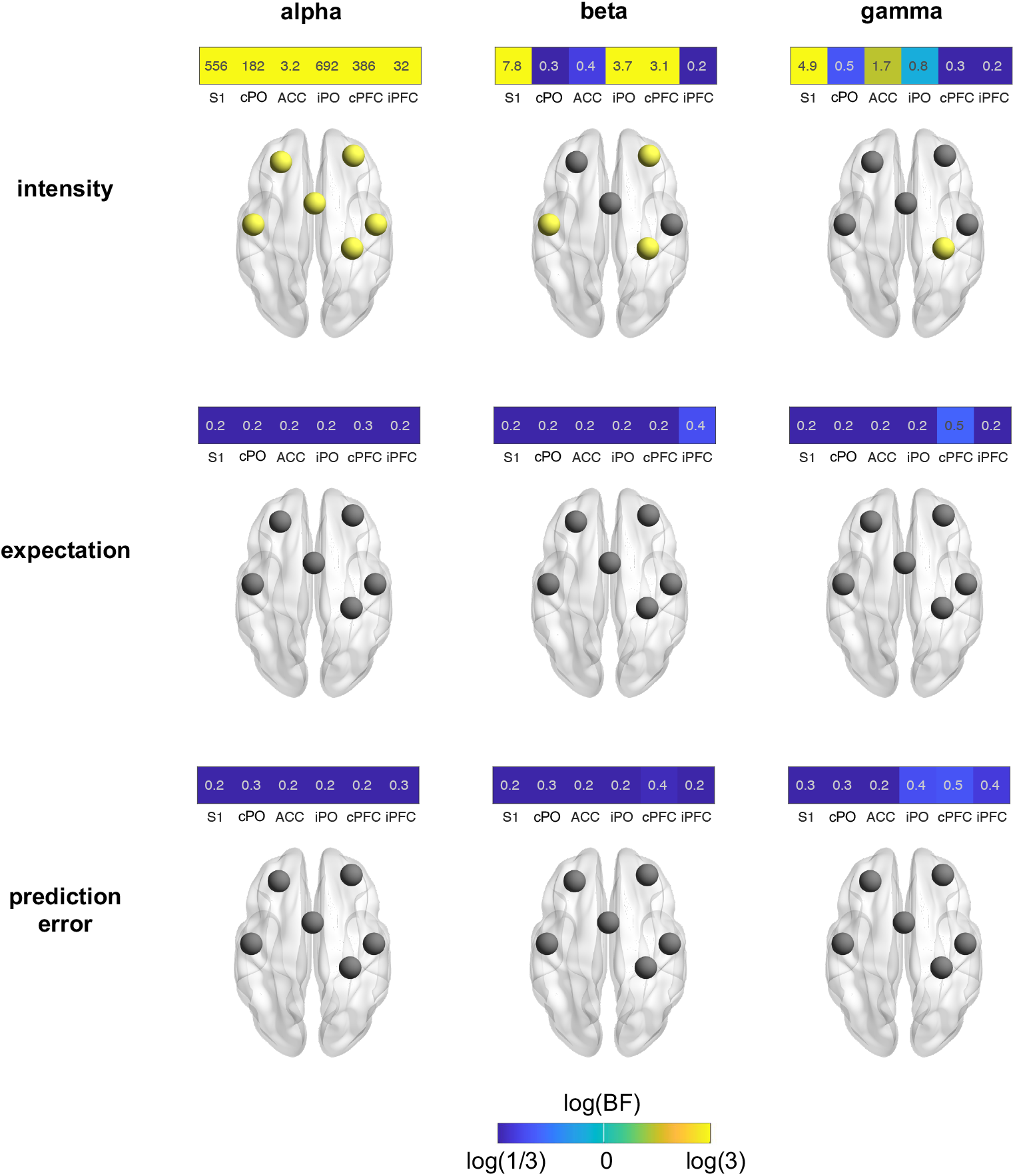
Effects of stimulus intensity, expectations, and prediction errors on local brain activity. Effects were assessed by Bayesian rmANOVAs with the factors intensity and expectation. The color of the heat map tiles scales with the log of the Bayes factor. It ranges from blue (BF < 1/3, at least moderate evidence against an effect) to yellow (BF > 3, at least moderate evidence for an effect). Brain images display ROIs in yellow which exhibit at least moderate evidence for an effect (BF > 3).

In contrast, we found weak to moderate evidence against effects of expectations or PEs on local brain activity at all frequency bands. Control analyses using shorter time windows showed qualitatively similar results (Figure S2).

In summary, we found that stimulus intensity but not expectations or PEs significantly influenced local oscillatory brain activity in response to brief painful stimuli.

### The effects of stimulus intensity and expectations on inter-regional functional connectivity

We next investigated how stimulus intensity and expectations influenced communication in our core network associated with pain processing. We therefore determined pairwise inter-regional connectivity in a network of six ROIs resulting in 15 connectivity values. These analyses were performed separately for the alpha, beta, and gamma frequency bands in the 1 s post-stimulus interval. Figure 7 shows the results of Bayesian rmANOVAs.

**Figure 7.**
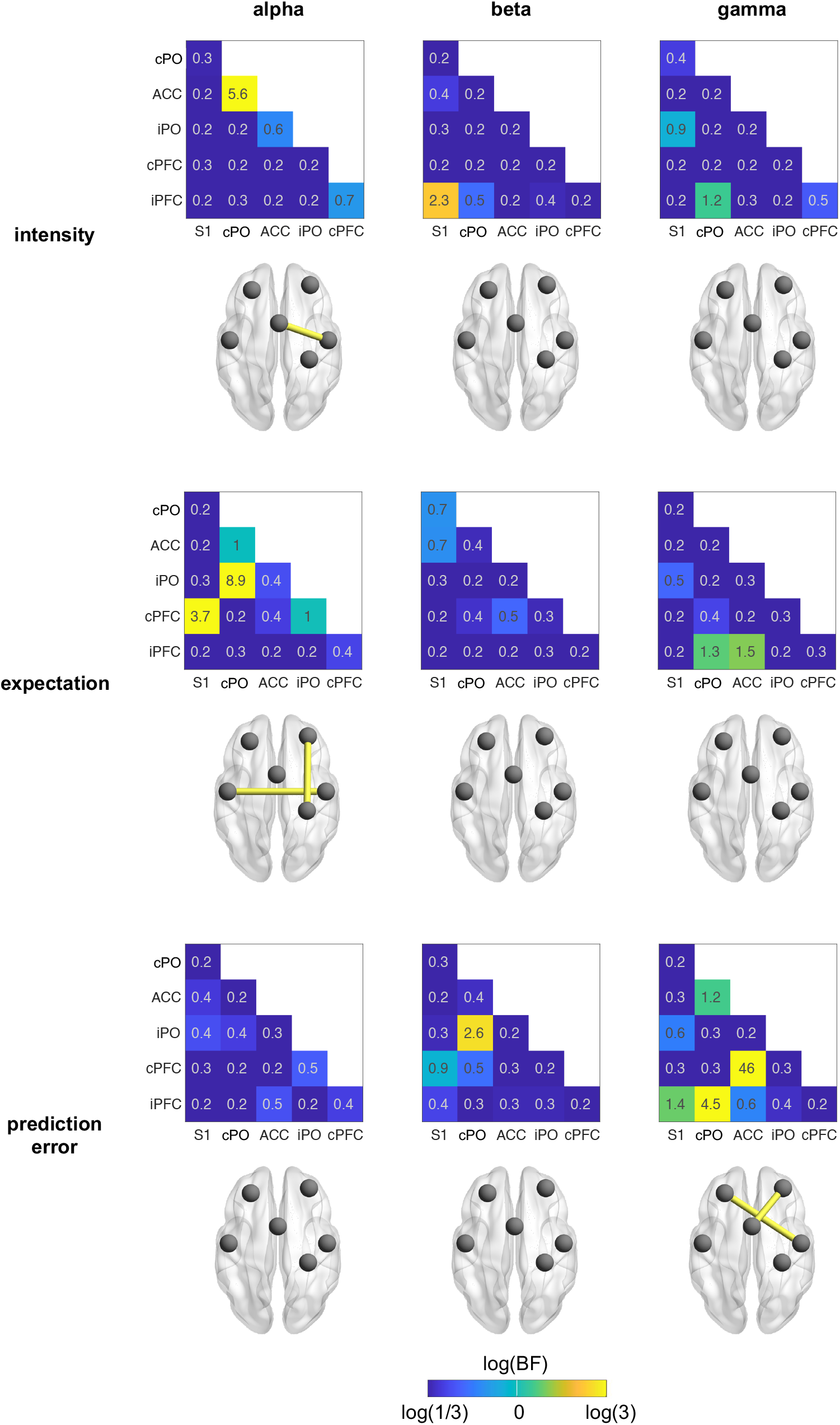
Effects of stimulus intensity, expectations, and prediction errors on inter-regional functional connectivity. Effects were assessed by Bayesian rmANOVA with the factors intensity and expectation. The color of the heat map tiles scales with the log of the Bayes factor. It ranges from blue (BF < 1/3, at least moderate evidence against an effect) to yellow (BF > 3, at least moderate evidence for an effect). Brain images display connections in yellow which exhibit at least moderate evidence for an effect (BF > 3).

We found moderate evidence for a stimulus intensity effect on the cPO – ACC connection in the alpha band. Here, connectivity was higher in the hi than the li condition. For most other connections and frequency bands, we found weak to moderate evidence against stimulus intensity effects.

Effects of expectation were found in the alpha band exclusively. We specifically observed moderate evidence for an expectation effect on the cPFC – S1 and iPO – cPO connections. In these connections, connectivity was lower in the HE than the LE conditions. For most other connections and frequency bands, we found weak to moderate evidence against expectation effects.

PE effects were observed in the gamma band exclusively. We found moderate to strong evidence for a PE effect on the cPFC – ACC and iPFC – PO connections. Specifically, the mean connectivity values of mismatch conditions (hiLE, liHE) were lower than those of non-mismatch conditions (liLE, hiHE). In other words, conditions involving a PE exhibited lower connectivity than those without a PE. For most other connections and frequencies, we observed weak to moderate evidence against a PE effect.

Taken together, we found that stimulus intensity and expectation influenced connectivity at alpha frequencies whereas PE effects were found at gamma frequencies.

### Direction of functional connectivity

The previous analyses showed that stimulus intensity, expectations and PEs modulated functional connectivity at alpha and gamma frequencies in a core network associated with pain processing. We were next interested to assess the direction of information flow for connections in which we found at least moderate evidence for intensity, expectation, and/or PE effects. To this end, we calculated an asymmetry score of directed connectivity between pairs of brain regions. The score was based on the bivariate partial directed coherence (PDC, (*38*)) measure and ranged from −1 to 1. The absolute value and the sign of the score indicate the strength and the direction of asymmetry, respectively. For the cPO-ACC connection, for which intensity effects were observed in the alpha band, we found strong evidence (BF = 13.4) that information flowed from cPO to ACC. For the cPFC-S1 connection, for which expectation effects were observed in the alpha band, we found strong evidence (BF = 13.1) that information flowed from cPFC to S1. For the other connections and frequency bands, we did not find evidence for an asymmetry of information flow. Thus, as summarized in Figure 8, for connections showing intensity effects, we found information flow predominantly from sensory to higher-order brain areas. Conversely, for connections displaying expectation effects, we found information flow predominantly from higher-order to sensory brain areas.

**Figure 8.**
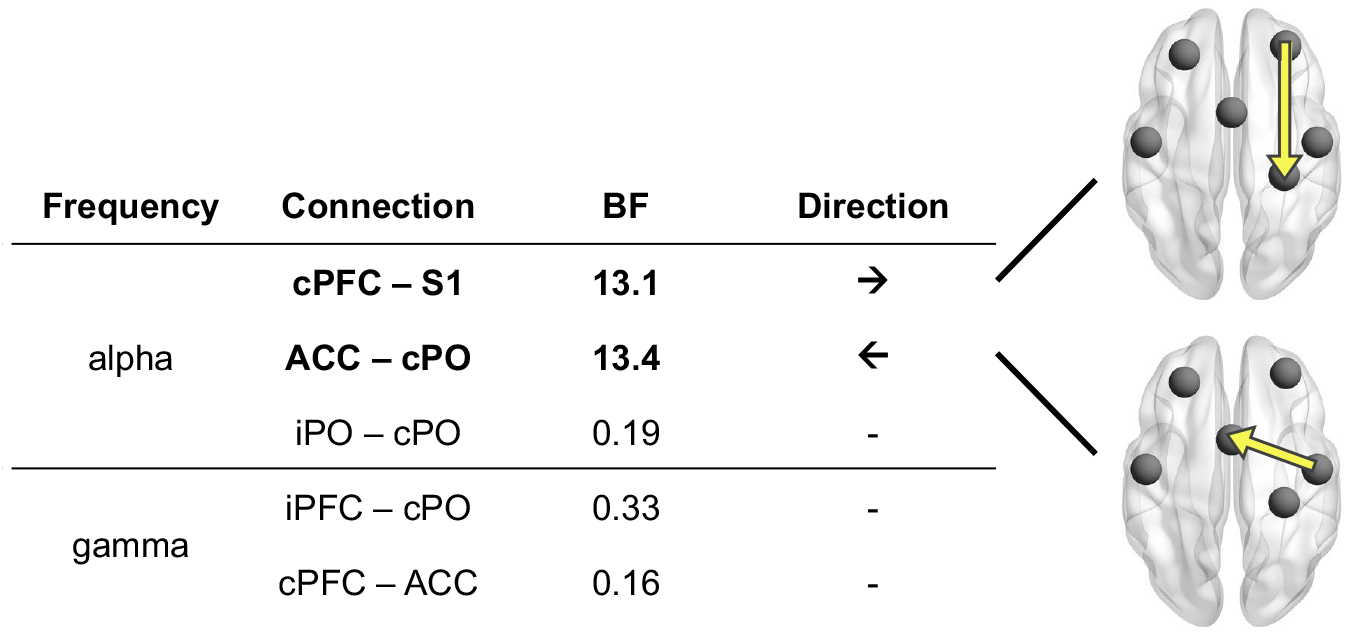
Direction of functional connectivity. Using an asymmetry score based on the PDC connectivity metric, we assessed the direction of information flow in connections which exhibited evidence for an effect in the previous connectivity analysis. Brain images depict connections with strong evidence for asymmetric information flow. The arrows indicate the dominant direction of information flow.

### Comparisons of stimulus intensity and expectations effects on local oscillatory brain activity and inter-regional functional connectivity

The previous analyses indicated that local brain activity and inter-regional connectivity differentially serve sensory and expectation effects on pain. We specifically observed that stimulus intensity shaped local brain activity more than inter-regional connectivity, while expectations and PEs shaped inter-regional connectivity more than local activity. To address this statistically, we conducted a Bayesian comparison of activity and connectivity models predicting the levels of stimulus intensity, expectation and PE. We found decisive evidence that activity models predicted stimulus intensity better than connectivity models (BF_pow/conn_ > 10^5^). Conversely, there was decisive evidence that connectivity models predicted expectations (BF_conn/pow_ > 10^2^) and PEs (BF_conn/pow_ > 2*10^2^) better than activity models.

### Summary

Figure 9 summarizes the main findings. On the behavioral level, both stimulus intensity and expectation significantly modulated the perception of pain. As expected, both higher stimulus intensities and expectations of stronger stimuli evoked higher pain ratings. In the brain, stimulus intensity effects were predominantly associated with changes of local brain activity. Stronger stimuli yielded stronger responses to brief painful stimuli in alpha, beta, and gamma frequency bands. In contrast, expectation effects on pain were associated with changes of inter-regional functional connectivity but not with changes of local brain activity. We particularly found that expectation effects were associated with top-down connectivity at alpha frequencies from cPFC to S1 and with connectivity between cPO and iPO. PEs were associated with changes of gamma-band connectivity exclusively. Bayesian model comparisons confirmed the differential involvement of local activity and inter-regional connectivity in sensory and expectation effects on pain. Specifically, stimulus intensity has a stronger influence on local brain activity than on inter-regional connectivity. Vice versa, expectations and PEs shape inter-regional connectivity more than local brain activity.

**Figure 9.**
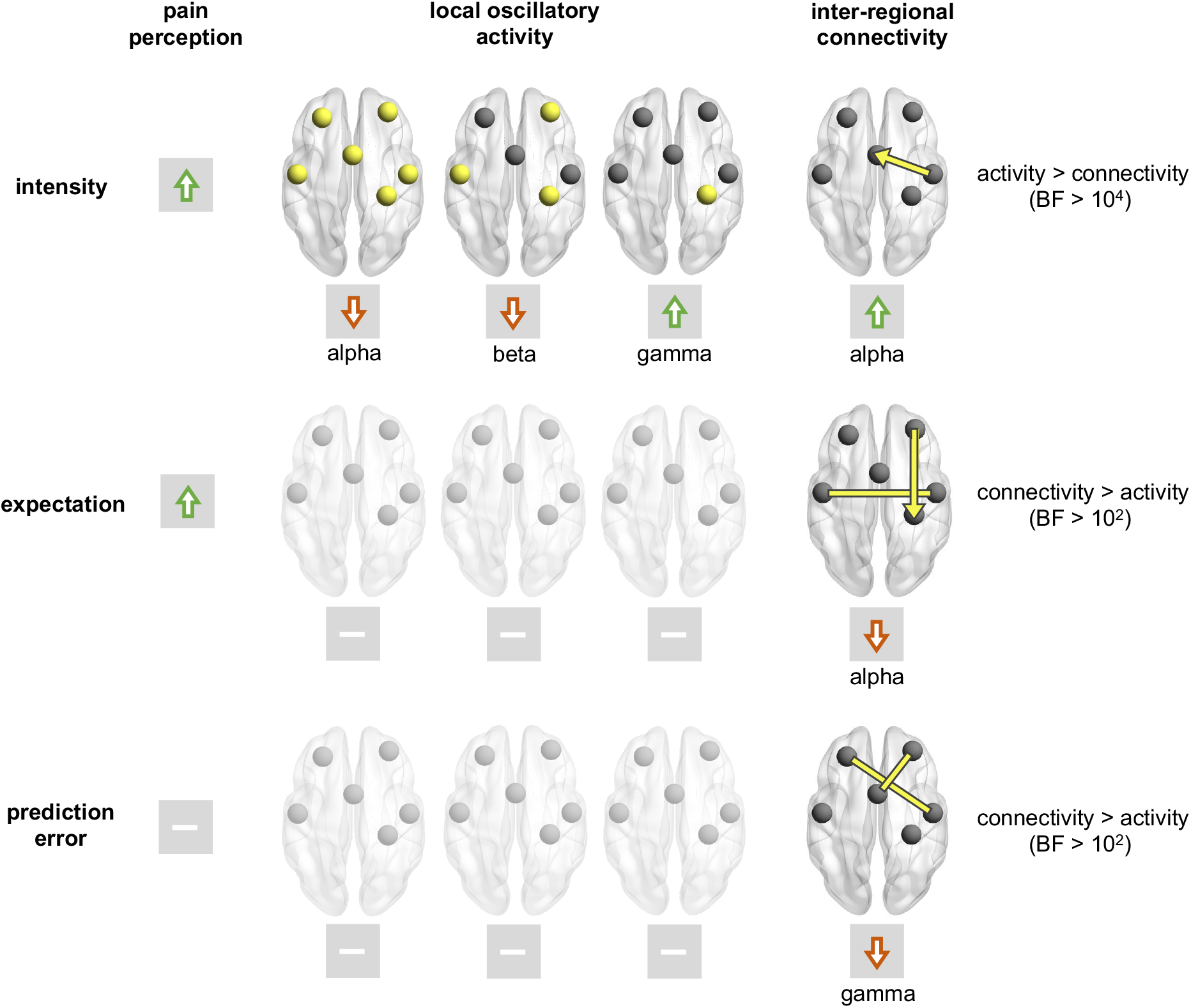
Synopsis of the effects of stimulus intensity, expectations, and prediction errors on pain perception, local brain activity, and inter-regional functional connectivity. Increases of stimulus intensity led to increases of pain ratings and local brain activity at gamma frequencies as well as to decreases of brain activity at alpha and beta frequencies. Expectations of stronger pain yielded increases of pain ratings and reduced connectivity between cPO and iPO and from cPFC to S1 at alpha frequencies. In contrast, expectations did not modulate local brain activity at any ROI and any frequency band. PEs did not change pain ratings or local brain activity but iPFC-cPO and cPFC-ACC connectivity at gamma frequencies. The last column shows the results of a Bayesian comparison of local brain activity and connectivity models predicting intensity, expectation and prediction errors.

## Discussion

In the present study, we investigated how the brain serves sensory and contextual effects on pain. To this end, we applied noxious stimuli to healthy human participants and independently modulated stimulus intensity and expectations. Pain ratings confirmed that stimulus intensity and expectation both influenced pain perception. Analyses of EEG recordings revealed that sensory and expectation effects on pain were served by fundamentally different brain mechanisms. In a core network associated with the processing of pain, sensory information shaped local oscillatory brain activity rather than inter-regional functional connectivity. In contrast, expectation and prediction errors influenced inter-regional functional connectivity but not local oscillatory brain activity.

### Sensory and expectation effects on local oscillatory brain activity and inter-regional functional connectivity

We observed that sensory information shapes local oscillatory brain activity significantly more than inter-regional connectivity. The effects of stimulus intensity on local oscillatory activity in various frequency bands are in accordance with previous EEG and MEG studies (*24, 33, 43, 44*). However, the effects of stimulus intensity on local brain activity and inter-regional connectivity have not been directly compared so far.

We further observed that expectations influenced inter-regional functional connectivity but not local oscillatory brain activity. To the best of our knowledge, expectation effects on functional connectivity have not yet been investigated by neurophysiological recordings. A few studies have investigated expectation effects on local oscillatory brain activity (*24, 25, 45, 46*). Their findings were inconsistent. Some studies found that expectations of high pain were associated with increased alpha activity (*24, 45*), others report unchanged (*25*), or decreased alpha activity (*46*). The present findings do not rule out any expectation effects on local brain activity. However, the crucial finding here is not the lack of expectation effects on local oscillatory activity, but that expectation effects on connectivity are significantly stronger than on local oscillatory activity.

### Expectation and prediction error signaling in the processing of pain

We found that expectation and prediction errors influenced connectivity at alpha/beta and gamma frequencies, respectively. This observation can be interpreted with reference to predictive coding (PC) frameworks of brain function. PC is a general theory used to explain how perception arises from the integration of sensory information and expectations (*26*). The framework proposes that the brain maintains an internal model of the environment which continuously generates predictions about sensory input. Discrepancies between these predictions and the actual sensory evidence, i.e. PEs, serve to adjust the internal model. In this way, the brain allocates its limited resources to events that are behaviorally relevant and useful for updating predictions, i.e., learning. It has been suggested that alpha and beta oscillations serve the signaling of predictions, whereas gamma oscillations have been proposed to signal PEs (*28–30, 39*). The present findings are in good accordance with this framework. They specify that expectation effects on pain might be particularly related to connectivity at alpha frequencies from the prefrontal to the somatosensory cortex. Specifically, expecting less pain was associated with relatively stronger connectivity. This implies that alpha-band connectivity might be mechanistically involved in an active down-regulation of nociceptive input. Prediction errors on the other hand were reflected in reduced gamma connectivity indicating that they are signaled in the brain in terms of a disruption of inter-regional communication which is in line with a recent study on PE signaling in the processing of pain (*24*).

### Distinct brain mechanisms serve sensory and expectation effects on pain

The key finding of our study is that sensory and expectation effects on pain are served by distinct brain mechanisms. Previous fMRI studies have already revealed that sensory and contextual effects on pain are associated with different spatial patterns of brain activity. For instance, one spatial pattern of brain activity termed the neurologic pain signature (NPS) is much more sensitive to sensory than to contextual effects on pain (*47, 48*). Vice versa, another pattern of brain activity termed the stimulus intensity independent pain signature (SIIPS) is sensitive to contextual but not to sensory effects on pain (*22*). Moreover, a spatial dissociation between the encoding of sensory information and expectations has also been found within the insular cortex (*39*).

Our results extend these findings by showing that not only the spatial brain activity patterns serving sensory and contextual effects on pain differ but that these effects are served by fundamentally different neurophysiological mechanisms. Sensory effects predominantly occurred in local brain oscillations whereas expectation effects were exclusively observed in inter-regional connectivity. The dissociation of sensory and expectation effects suggests that both physiological phenomena are partially independent of each other.

These findings might have implications for the understanding, assessment, and treatment of clinical pain conditions. In acute pain, which is predominantly shaped by sensory information, assessing and modulating local oscillatory brain activity might be appropriate. In contrast, in chronic pain, which is often largely detached from sensory information, inter-regional connectivity might be more informative than local activity. Such a close association between inter-regional connectivity and chronic pain is in accordance with studies using fMRI (*15, 49, 50*) and recent EEG studies on connectivity in chronic pain (*51, 52*) and psychiatric disorders (*53*). In this way, the present findings can help to guide the development of biomarkers of acute and chronic pain. Beyond, our results might inform the search for neuronal targets for invasive as well as non-invasive interventions aiming at alleviating pain.

### Limitations

When interpreting our findings, certain limitations should be considered. First, in our paradigm, the effects of expectations on pain perception were weaker than the effects of stimulus intensity. The lack of expectation effects on local brain activity might therefore reflect the weak expectation effects on pain perception, and other paradigms with stronger expectation effects on perception might well modulate local brain activity. However, the central finding of the present study is not the absolute strength of sensory and expectation effects but that the patterns of sensory and expectation effects on local brain oscillations and brain connectivity fundamentally differ. The strength of perceptual effects might well determine the strength of neurophysiological effects but is unlikely to fundamentally change the difference in the patterns of sensory and expectation effects on brain activity and connectivity. We are therefore confident that the present findings reflect a fundamental difference in the brain mechanisms serving sensory and expectation effects on pain.

Second, to modulate pain, we manipulated participants’ expectations. Expectations are a particularly powerful and clinically highly relevant modulator of pain (*4–7*). However, it is unclear whether the present observations are specific to expectation-induced modulations of pain or whether they generalize to other cognitive and contextual modulations of pain.

Third, we applied brief experimental pain stimuli to healthy human participants. It is unclear whether these findings can be translated to other experimental and clinical types of pain. It remains to be investigated whether the findings generalize to chronic pain conditions in which other brain mechanisms apply and in which the brain undergoes substantial structural and functional plasticity (*11, 54*).

### Conclusions

Taken together, the present study shows that sensory and expectation effects on pain are served by distinct brain mechanisms. Sensory effects on pain are served by changes of local oscillatory brain activity, whereas expectation effects and discrepancies between sensory information and expectations are served by changes of inter-regional functional connectivity. These results provide novel basic science insights into the brain mechanisms of pain and analgesia. They specifically advance the understanding of how the brain serves key modulations of the subjective experience of pain. Beyond, they can inform the development of novel tools for the assessment and treatment of clinical pain conditions.

## Materials and methods

### Participants

The study was performed in healthy human participants at the university hospital of the Technical University of Munich (TUM). Written informed consent was obtained from all participants prior to the experiment. The Ethics Committee of the Medical Faculty of the TUM approved the study protocol. The study was preregistered at ClinicalTrials.gov (NCT04296968) and conducted in accordance with the latest version of the Declaration of Helsinki. It followed recent guidelines for the analysis and sharing of EEG data (*55*). Inclusion criteria were right-handedness and age >18 years. Exclusion criteria were pregnancy, neurological or psychiatric diseases, and regular intake of medication (aside from contraception and thyroidal medication). Severe internal diseases (e.g. diabetes) and skin diseases (e.g. psoriasis, vitiligo), previous surgeries at the head or spine, current or recurrent pain, metal or electronic implants, and any previous side effects associated with thermal stimulation constituted additional exclusion criteria.

For the current rmANOVA design (one group, four measurements), an assessment of statistical power using G*Power (*56*) yielded a sample size estimate of 36 participants with a power of 0.95, an alpha of 0.05, and medium effect sizes of f = 0.25 (corresponding to an η^2^ of 0.06 (*57*)).

The original study recruited 58 healthy human participants (29 females, age: 24.0 ± 4.3 y [mean ± SD]). Ten participants were excluded due to either the absence of pain or low pain ratings [<10 on a numerical rating scale from 0 (no pain) to 100 (maximum tolerable pain)] during the familiarization run (n = 8), excessive startle responses in response to painful stimulation during the training run (n = 1), or technical issues with the response box used during catch trials (n = 1). To ensure robust estimates of connectivity values, we here additionally excluded participants with less than 10 trials remaining after the raw data cleaning procedure described below (n = 8). The final data set used here thus comprised 40 participants (all right-handed, 21 females, age: 23.4 ± 2.9 y). Average anxiety and depression scores were below clinically relevant cutoff scores of 8/21 (*58*) on the Hospital Anxiety and Depression Scale (*59*) (anxiety: 3.2 ± 2.2; depression: 0.9 ± 1.2).

### Procedure

The objective of this analysis was to assess how sensory and contextual modulations are served by local brain activity and inter-regional brain connectivity. The experiment involved two levels of noxious stimulus intensities (hi and li) and two types of visual cues (HE and LE) resulting in four experimental conditions. The visual cues probabilistically predicted the intensity of the subsequent noxious stimulus. The high expectation (HE) cue was followed by a hi stimulus in 75% of the trials and by a li stimulus in 25% of the trials. Vice versa, the low expectation (LE) cue was followed by a hi stimulus in 25% of the trials and by a li stimulus in 75% of the trials (Figure **1**a).

Figure **1**b depicts the sequence of events for each trial. After a variable fixation period ranging from 1.5 to 3 s, a visual cue (either blue dot or yellow square) was displayed for 1 s. A brief painful heat stimulus was applied 1.5 s after cue offset. 3 s after the painful stimulus, participants were visually prompted to provide a verbal rating of the perceived pain intensity on a numerical rating scale ranging from 0 (no pain) to 100 (maximum tolerable pain in the context of the experiment). To ensure that participants continuously paid attention to the visual cues, participants were visually prompted to indicate by a button press whether a HE or a LE cue had been presented last in 10% of the trials. An average accuracy of 95.6 ± 0.1% indicated that participants successfully focused on the task during the entire experiment. Trials were separated by a 3 s period during which a white fixation cross was presented.

The experiment consisted of four runs with 40 trials each (hiHE [n = 15], hiLE [n = 5], liLE [n = 15], liHE [n = 5]), resulting in total trial numbers of hiHE [n = 60], hiLE [n = 20], liLE [n = 60], liHE [n = 20]. Runs were separated by short breaks of ~3 min. Pairings of visual cues with stimulus intensities were balanced across participants.

Prior to the experiment, the participants were familiarized with the stimulation and the intensity rating procedure by applying a sequence of 10 heat stimuli. Next, participants were informed about the pairing between cues and stimulus intensities and a training run comprising 16 trials was conducted. This was to ascertain that all participants were aware of the pairing and to minimize learning during the main experiment. During the experiment, participants sat in a comfortable chair. They wore protective goggles and listened to white noise on headphones to eliminate effects of ambient sounds. Please see (*23*) for additional details.

### Stimulation

A laser pulse with a wavelength of 1,340 nm, a duration of 4 ms and spot diameter of approximately 7 mm was used to apply painful stimuli to the left hand (*60*). For li and hi stimuli, the stimulus intensity was set to 3 and 3.5 J, respectively. These stimulus intensities are known to consistently elicit painful sensations of discriminable intensity (*60*). The stimulation site was slightly changed after each stimulus to avoid tissue damage and habituation or sensitization.

### Recordings and preprocessing

Brain activity was recorded using actiCAP snap/ slim with 64 active sensors (Easycap) placed according to the extended 10-20 system and BrainAmp MR plus amplifiers (Brain Products, Munich, Germany). During the recording, sensors were referenced to FCz and grounded at Fpz. The signals were sampled at 1,000 Hz (0.1-μV resolution) and band-pass filtered between 0.016 and 250 Hz while impedances were kept below 20 kΩ.

The BrainVision Analyzer software (version 2.1.1.327, Brain Products, Munich, Germany) was used for preprocessing. First, raw signals were low-pass filtered with a cutoff frequency of 225 Hz. After down-sampling to a rate of 500 Hz, a 1 Hz high-pass filter (fourth-order Butterworth) and a band-stop filter between 49 and 51 Hz filter removing line noise were applied. An independent component (IC) analysis based on the extended infomax algorithm was then conducted based on the −4.2 to 3.2 s peristimulus time windows of the EEG data. Subsequently, ICs representing artifacts originating from eye movements or muscles were removed from the unfiltered EEG data (*61*) using visual inspection. Moreover, data segments of 400 ms centered around data samples with amplitudes exceeding ±100 μV and data jumps exceeding 30 μV were automatically marked for rejection. Finally, the data were inspected visually and remaining artifacts were manually marked for rejection. All signals were re-referenced to the average reference. The cleaned data were exported to Matlab (version R2019b, Mathworks, Natick, MA) and further analyses were performed using FieldTrip [version 20210411 (*62*)]. Data were segmented into epochs ranging from −4 to 3 s in peristimulus time and all trials with marked artifacts or pain ratings of zero were excluded. This resulted in 49.5 ± 8.5, 16.8 ± 2.8, 18.0 ±1.6, and 52.9 ± 4.2 trials per participant in the liLE, liHE, hiLE, and hiHE conditions, respectively. To assure that all analyses for the different trial types were eventually performed on the same number of trials, we matched the numbers of trials. Figure S1 shows details of the trial matching procedure.

### Source model

To project sensor-level time series to source level, we employed Linearly Constrained Minimum Variance (LCMV) beamformers (*63*) implemented in FieldTrip (*62*). Frequency-specific array-gain LCMV spatial filters for alpha, beta and gamma frequencies were constructed based on a lead field and a frequency-specific covariance matrix. A boundary element approximation of a realistically shaped, three-shell head model was used as the lead field. For each individual and frequency band, the covariance matrix was computed from the band-pass filtered, −1 s to 1 s (peristimulus time) concatenated data segments of all (non-rejected) trials. To ensure a robust computation of the inverse of the covariance matrix we employed Tikhonov regularization as implemented in FieldTrip with a regularization parameter value of 5% of the average sensor power. The fixed orientation of the lead field for every source location was chosen to maximize the spatial filter output. Source-level signals were then obtained by applying the frequency-specific LCMV operator to the corresponding band-pass filtered sensor-level time series.

### Assessment of source-level time-frequency representations

Source-level time-frequency representations were obtained using the following procedure: First, we projected the band-pass filtered sensor-level signals to source space using five frequency-specific LCMV spatial filters (i.e., for frequencies <8 Hz, 8-12 Hz, 13-30 Hz, 30-60 Hz, and 60-100 Hz). For each ROI, we generated TFRs as well as time-courses of alpha, beta, and gamma brain activity. The TFRs are based on Hanning-tapered data. Time courses of brain activity were computed based on moving time windows and using a Slepian multi-taper approach (see below). TFRs and time-courses of brain activity were computed from data segments with widths of 500 ms and 250 ms for frequencies below and above 30 Hz, respectively. Both TFRs and time-courses of brain activity are displayed as percentage change relative to a baseline period ranging from 0.75 to 0.25 s before the stimulus. To maximize the signal-to-noise ratio for visualization, the results represent the grand average across participants and hi trials.

### Analysis of local brain activity

Local oscillatory brain activity was assessed as frequency-specific source power of the 6 ROIs. First, source level timeseries band-pass filtered to the frequency band of interest were obtained using the beamformer described above. For these signals, we computed the power of the frequency in the middle of the frequency band of interest using a Slepian multi-taper approach (*64*). The spectral smoothing width was set to one half of the width of the frequency band of interest. In this way, the power value incorporates information of the entire frequency band of interest. We computed source power in the alpha (8-12 Hz), beta (14-30 Hz) and gamma (60-100 Hz) frequency bands for each trial. We then averaged power values across trials for each condition and subject. To allow for the comparison of the effects on local brain activity to those on brain connectivity, the analysis was primarily performed on a 1 s post-stimulus interval. However, sensor-level findings indicate that the effects of painful stimuli on oscillatory brain activity are usually confined to shorter time windows. Specifically, pain-induced suppressions of brain activity at alpha and beta frequencies occur at latencies between 500 and 900 ms and between 300 and 600 ms, respectively (*33, 43*). In addition, pain-induced increases of brain activity at gamma frequencies occur between 150 and 350 ms (*44*). We therefore performed control analyses using these shorter time intervals (see Supplementary Figure S2 for results).

### Analysis of inter-regional connectivity

Connectivity analyses were performed on the 1 s post-stimulus intervals of the source level timeseries of the 6 ROIs. First, we computed the source level cross-spectral density of each participant using a multi-taper approach analogous to the one used for the computation of source power.

To assess functional connectivity, we calculated the debiased weighted Phase Lag Index (dwPLI, (*32*)) based on all trials of each condition and for every subject. We selected the dwPLI measure due to its insensitivity to volume conduction effects.

For the assessment of the direction of connectivity, we used an asymmetry score based on bivariate partial directed coherence (PDC, (*38*)). Specifically, for two ROIs A and B, the bivariate PDC analysis yields two values, PDC_A→B_ and PDC_B→A_, representing the directed connectivity strength from A to B and from B to A, respectively. We cast these two values into a single asymmetry score, (PDC_A→B_ - PDC_B→A_)/(PDC_A→B_ + PDC_B→A_), ranging from −1 to 1. A large absolute value of the asymmetry score indicates a strong asymmetry of directed connectivity. The sign of the asymmetry score reveals the predominant direction of information flow. Direction of connectivity was calculated for connections that had shown intensity, expectation, and/or PE effects in previous analyses. For connections with evidence for an intensity or expectation effect in the Bayesian ANOVA, we included all trial types in the computation of the asymmetry score. For connections with evidence for an interaction effect, we included trials with a mismatch between cue and intensity only.

### Statistical analyses

For each of the four trial types (liLE, hiLE, liHE, hiHE), behavioral and EEG measures were computed based on an identical number of trials. This number was determined as the minimum number of available trials across the four trial types. Details of the trial matching procedure can be found in the supplementary material (Figure S1).

Building upon previous investigations (*39, 40*), we made specific predictions about how EEG responses signaling stimulus intensity, expectations, PEs, or combinations thereof are modulated across the four trial types. To formally test these predictions, we performed repeated measures ANOVAs (rmANOVAs) with the independent variables stimulus intensity and expectation. In these rmANOVAs, responses signaling stimulus intensity and expectations would manifest as main effects, whereas responses signaling PEs would manifest as interactions. To quantify effects and to facilitate interpretation of negative findings, we performed Bayesian rmANOVAs (*41*).(*41*). In Bayesian rmANOVAs, the Bayes factor (BF) is the ratio between the likelihood of the data given the effect of interest and the likelihood of the data without the effect of interest. BF > 3 and BF > 10 indicate moderate and strong evidence in favor of the effect of interest, whereas BF < 1/3 and BF < 0.1 indicate moderate and strong evidence against the effect of interest, respectively (*41*). We considered a neural measure or pain rating as corresponding to the intensity or expectation pattern if there was at least moderate evidence for the corresponding main effect. Accordingly, we considered a neural measure or pain rating as corresponding to the prediction error pattern if the evidence for an interaction effect of intensity and expectation was at least moderate.

Lastly, for the assessment of asymmetry of information flow we tested asymmetry scores against 0 using a nonparametric Bayesian t-test.

All parametric Bayesian analyses were conducted using the BayesFactor package in R (*65*), for non-parametric Bayesian t-tests we used freely available R code (*66*).

### Bayesian model comparison

We intended to statistically assess whether an experimental contrast (intensity, expectation, or PE) is associated more strongly with local activity or inter-regional connectivity. To this end, we conducted a Bayesian comparison of power-based and connectivity-based models predicting the levels of intensity, expectation and PE. Specifically, we computed the Bayesian evidence of logistic models mapping individual power and connectivity values to the probability of observing a certain level of intensity, expectation, or PE. In the analysis, we consider N_pow_ = 6 power values and N_conn_ = 15 connectivity values in each of the N_freq_ = 3 frequency bands. For each of the three types of experimental contrasts, this resulted in N_freq_*N_pow_ = 18 model evidence values for the power-based models and N_freq_*N_conn_ = 45 model evidence values for the connectivity-based models. The Bayes factor for, e.g., the intensity manipulation reported in the manuscript is the average of the 18 power-based model evidence values divided by the average of the 45 connectivity-based model evidence values. For the factor expectation and the interaction between expectation and intensity, i.e., PE, we proceeded analogously. The derivation of Bayesian model comparisons for logistic regression models follows the description in (*67*) and is provided in the supplement.

### Data and code availability

All raw and preprocessed data in EEG-BIDS format are available at the open science framework (OSF, https://osf.io/jw8rv/). The code will be made available at OSF upon acceptance.

## Acknowledgments

We are thankful to Ulrike Bingel for valuable insights and comments on the manuscript.

## Funding

The study was supported by the Deutsche Forschungsgemeinschaft (PL 321/14-1) and the TUM Innovation Network *Neurotechnology in Mental Health* (Neurotech).

## Author contributions

- Conceptualization: FSB, MMN, MP
- Software: FSB
- Formal analysis: FSB
- Methodology: FSB, MP, JG
- Investigation: FSB, MMN
- Visualization: FSB, MP
- Supervision: MP, JG
- Writing — original draft: FSB, MP
- Writing — review & editing: FSB, MMN, VDH, ESM, CGA, LT, JG, MP
- Project administration: MP
- Funding acquisition: MP

## Competing interests

No competing interests to declare.

## Supplementary Information

### Trial matching

**Figure S1.**
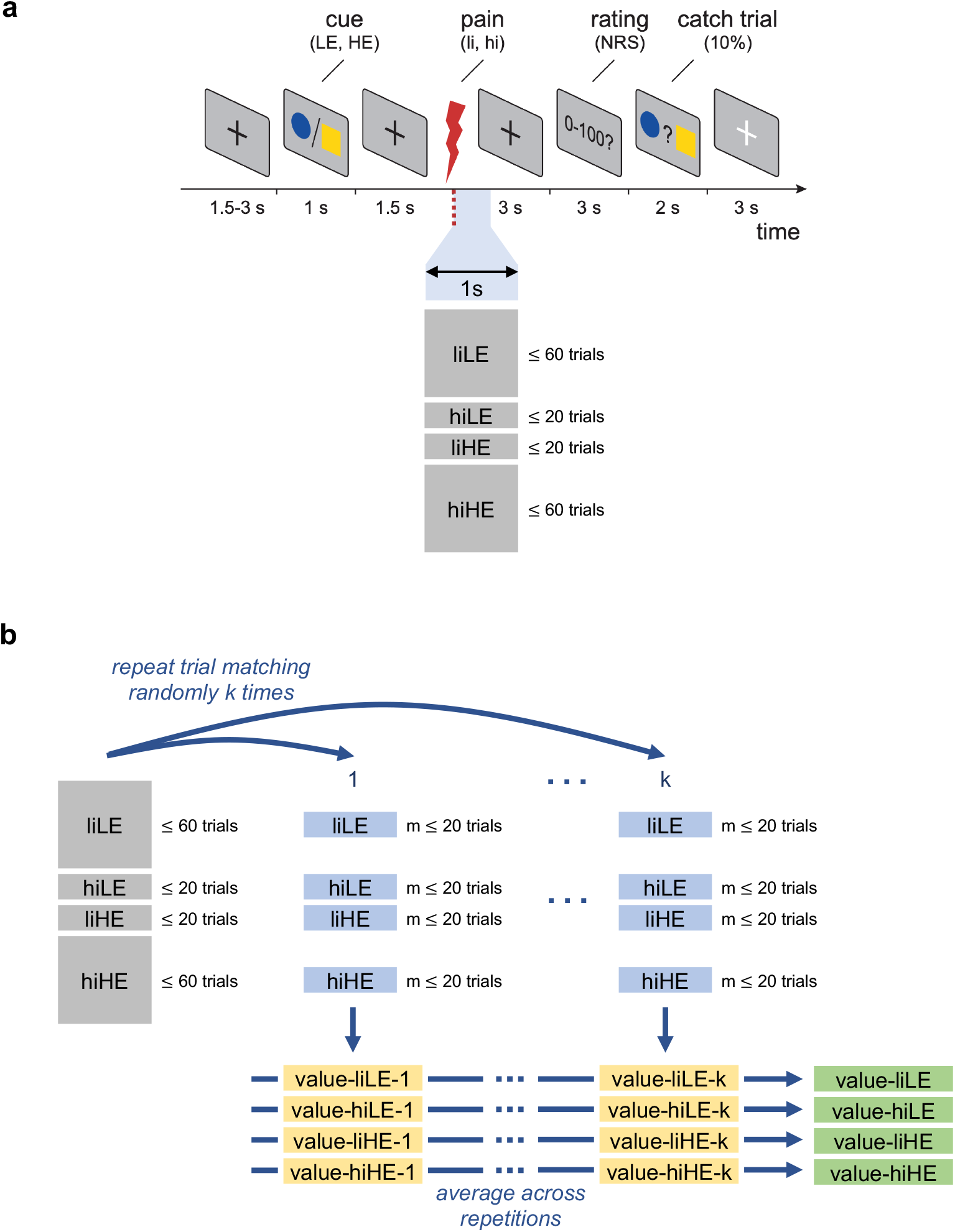
Trial matching. (a) The experiment comprised 160 trials per participant. In each trial, a cue (LE/HE) was presented which probabilistically predicted the intensity of a subsequent painful stimulus (li/hi). A LE(HE) cue preceded a li(hi) stimulus in 75% and a hi(li) stimulus in 25% of trials. This design in combination with the rejection of bad trials resulted in an unbalanced number of trials across the four trial types liLE, hiLE, liHE, and hiHE. (b) In order to circumvent a sample-size bias problem, all neural measures were computed based on the same number (m ≤ 20) of trials. The matching of trial sets was done randomly and repeated k = 256 times. For each trial set, the corresponding values (pain rating/power/connectivity) were computed. This resulted in k estimates per measure which were averaged to obtain a single value per measure.

### The effects of stimulus intensity and expectations on local brain activity in time windows showing strongest responses

**Figure S2.**
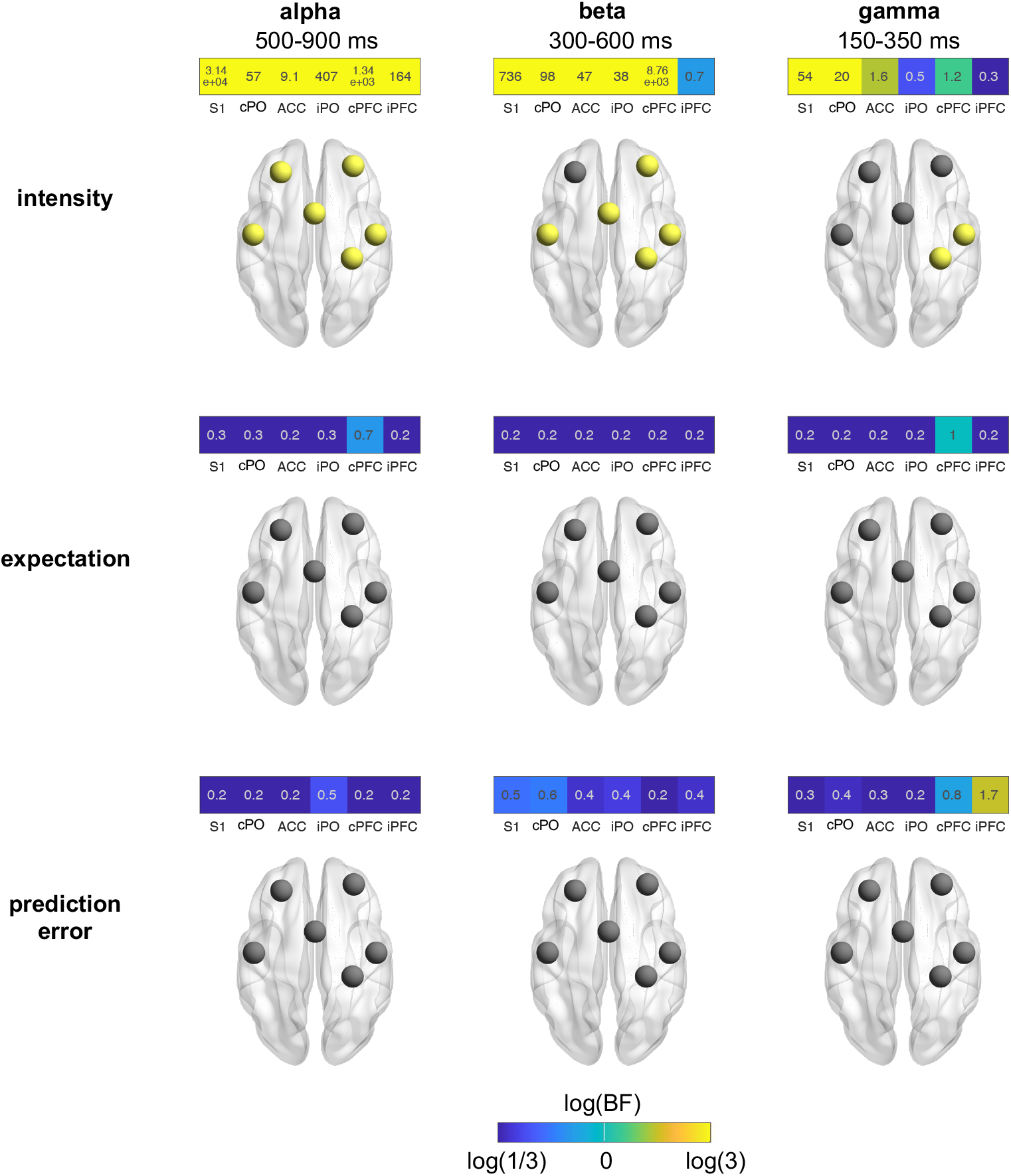
Effects of stimulus intensity, expectations, and prediction errors on local brain activity. Power at alpha, beta, and gamma frequencies was quantified using the time windows 500-900 ms, 300-600 ms, and 150-350 ms, respectively. Heat maps indicate Bayes factors of a Bayesian rmANOVA with factors intensity and expectation. The color of the heat map tiles scales with the log of the Bayes factor. It ranges from blue (BF < 1/3, at least moderate evidence against an effect) to yellow (BF > 3, at least moderate evidence for an effect). Brain schematics display ROIs in yellow which exhibit at least moderate evidence for an effect (BF > 3).

### Bayesian model comparison

The objective is to statistically assess whether an experimental contrast (intensity, expectation, or PE) is associated more strongly with regional activity or inter-regional connectivity. To this end, we conducted a Bayesian comparion of power-based and connectivity-based models predicting the levels of intensity, expectation, and PE. The derivations in this section follow the description in Bishop et al. (2006).

Say there are *N* participants to be included in the analysis. Let **S**^*i*^, **E**^*i*^, **P**^*i*^, and **C**^*i*^ be the data vectors associated with participant *i* ∈ 1, …, *N*. The vectors **S**^*i*^ = [0, 0, 1, 1]^⊤^ and **E**^*i*^ ∈ [0, 1, 0, 1]^⊤^ encode the levels of stimulus intensity (li: 0, hi: 1) and expectation (LE: 0, HE: 1), respectively. The vectors 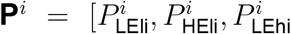 and 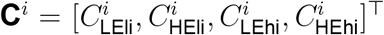 contain the corresponding values of power and connectivity, respectively. In the following, we will solely focus on the comparison of models discriminating low and high intensity. The derivation of the comparison of models for the expectation and PE contrasts is analogous.

First, to arrive at a single binary dependent and a single continuous independent variable (per model), we average the data across the two levels of expectation. In addition, to account for the repeated measures design of the experiment, we center the participant-level independent variable at 0. Thus, formally:

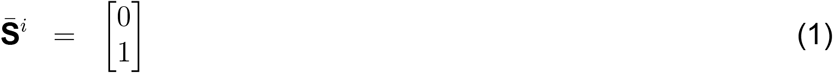

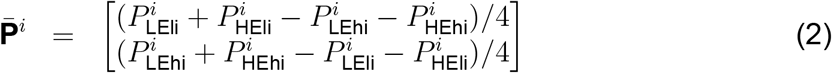

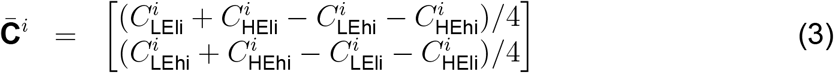

The data of all participants is cobined in vectors 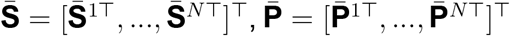, and 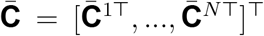. To be able to use the same prior for all independent variables, the data vectors of the independent variables are scaled by their standard deviation:

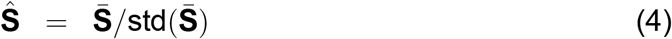

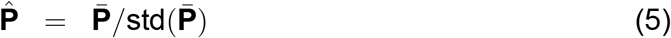

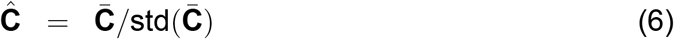

In both models to be compared, the probabitliy of observing the high intensity (hi) level is modeled as a logistic function:

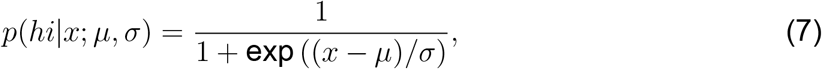

where *μ* and *σ* are parameters controlling the location and scale of the logistic function, respectively. Depending on the type of model, the continuous independent variable *x* represents either a power or a connectivity value. The likelihood of the data given the model parameters thus is

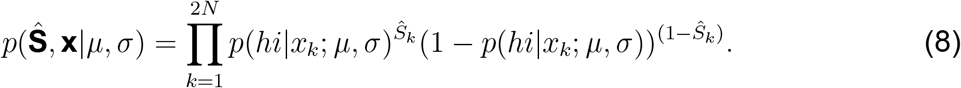

with

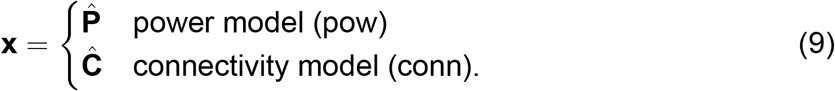

For the computation of the Bayesian model evidence a prior distirbution over the model paramters *μ* and *σ* must be specified. Here, we select a bivariate standard normal distribution:

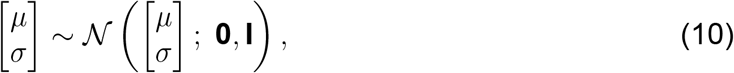

where **0** and **I** are the zero vector and identity matrix, respectively. Figure S3 shows graphs of logistic functions for several parameter values drawn from the prior distribution.

**Figure S3.**
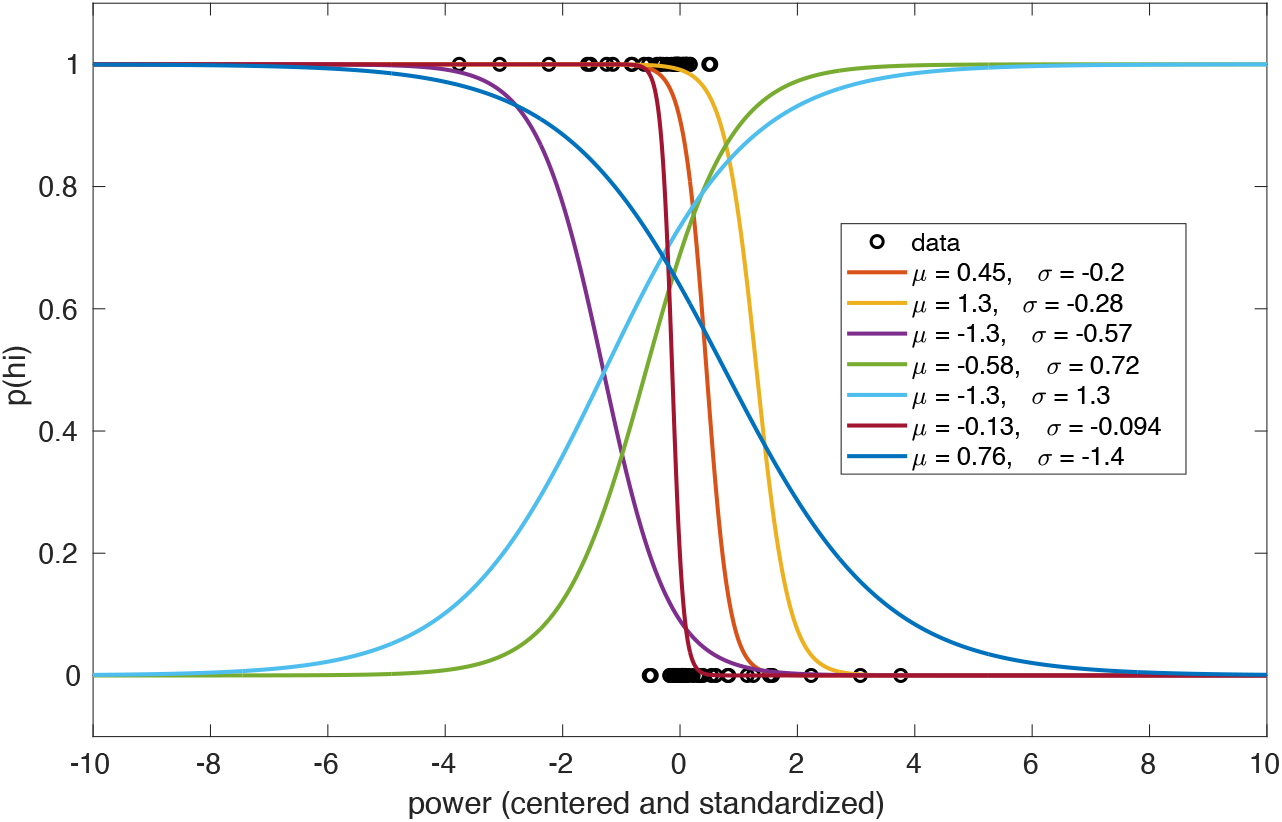
Graphs of logistic functions for several parameter values drawn from the prior distribution specified above. To show the prior graphs in relation to the data, by way of example, power values of ROI S1 are depicted as black circles. Specifically, the x- and y-values of the data points correspond to the values in vectors 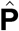 and 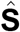, respectively.

According to Bayes’ rule, the probability of the data given the model (a.k.a. model evidence) is

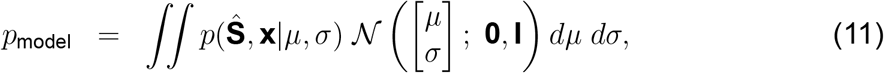

where for the power and connectivity models, **x** is substituted by 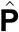 and 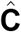, respectively. In our implementation, we compute this integral using standard Monte Carlo integration with 10^5^ samples.

The described procedure results in *N*_pow_ = 6 power and *N*_conn_ = 15 connectivity values per frequency band. The model evidence is computed for all individual power and connectivity values at all *N*_freq_ = 3 frequency bands. The resulting model evidence values are denoted by 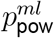 and 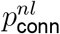 with indices *m, n*, and *l* coding for the different power, connectivity and frequency values, respectively. The Bayes factors reported in the manuscript represent the ratio of averaged model evidences:

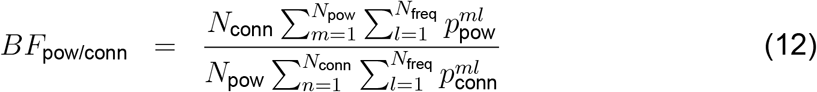

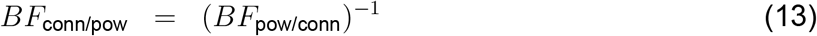

## Notes

### Competing Interest Statement

The authors have declared no competing interest.

## References

1. L. Y. Atlas, T. D. Wager, How expectations shape pain. Neurosci Lett 520, 140–148 (2012).

2. H. L. Fields, How expectations influence pain. Pain 159 Suppl 1, S3–S10 (2018).

3. K. J. Peerdeman, A. I. van Laarhoven, M. L. Peters, A. W. Evers, An Integrative Review of the Influence of Expectancies on Pain. Front Psychol 7, 1270 (2016).

4. P. Enck, U. Bingel, M. Schedlowski, W. Rief, The placebo response in medicine: minimize, maximize or personalize? Nat Rev Drug Discov 12, 191–204 (2013).

5. D. G. Finniss, T. J. Kaptchuk, F. Miller, F. Benedetti, Biological, clinical, and ethical advances of placebo effects. Lancet 375, 686–695 (2010).

6. T. D. Wager, L. Y. Atlas, The neuroscience of placebo effects: connecting context, learning and health. Nat Rev Neurosci 16, 403–418 (2015).

7. U. Bingel, Placebo 2.0: the impact of expectations on analgesic treatment outcome. Pain 161 Suppl 1, S48–S56 (2020).

8. J. W. Vlaeyen, G. Crombez, S. J. Linton, The fear-avoidance model of pain. Pain 157, 1588–1589 (2016).

9. S. Cormier, G. L. Lavigne, M. Choiniere, P. Rainville, Expectations predict chronic pain treatment outcomes. Pain 157, 329–338 (2016).

10. K. J. Peerdeman, A. I. van Laarhoven, S. M. Keij, L. Vase, M. M. Rovers, M. L. Peters, A. W. Evers, Relieving patients’ pain with expectation interventions: a meta-analysis. Pain 157, 1179–1191 (2016).

11. M. N. Baliki, A. V. Apkarian, Nociception, Pain, Negative Moods, and Behavior Selection. Neuron 87, 474–491 (2015).

12. L. Garcia-Larrea, R. Peyron, Pain matrices and neuropathic pain matrices: a review. Pain 154 Suppl 1, S29–43 (2013).

13. M. Ploner, C. Sorg, J. Gross, Brain Rhythms of Pain. Trends Cogn Sci 21, 100–110 (2017).

14. A. Kucyi, K. D. Davis, The dynamic pain connectome. Trends Neurosci 38, 86–95 (2015).

15. J. J. Lee, H. J. Kim, M. Ceko, B. Y. Park, S. A. Lee, H. Park, … C. W. Woo, A neuroimaging biomarker for sustained experimental and clinical pain. Nat Med 27, 174–182 (2021).

16. M. M. Nickel, S. Ta Dinh, E. S. May, L. Tiemann, V. D. Hohn, J. Gross, M. Ploner, Neural oscillations and connectivity characterizing the state of tonic experimental pain in humans. Hum Brain Mapp 41, 17–29 (2020).

17. T. Spisak, B. Kincses, F. Schlitt, M. Zunhammer, T. Schmidt-Wilcke, Z. T. Kincses, U. Bingel, Pain-free resting-state functional brain connectivity predicts individual pain sensitivity. Nat Commun 11, 187 (2020).

18. P. Taesler, M. Rose, Prestimulus Theta Oscillations and Connectivity Modulate Pain Perception. J Neurosci 36, 5026–5033 (2016).

19. P. Tetreault, A. Mansour, E. Vachon-Presseau, T. J. Schnitzer, A. V. Apkarian, M. N. Baliki, Brain Connectivity Predicts Placebo Response across Chronic Pain Clinical Trials. PLoS Biol 14, e1002570 (2016).

20. S. Vanneste, D. De Ridder, Chronic pain as a brain imbalance between pain input and pain suppression. Brain Communications 3, (2021).

21. T. D. Wager, L. Y. Atlas, M. A. Lindquist, M. Roy, C. W. Woo, E. Kross, An fMRI-based neurologic signature of physical pain. N Engl J Med 368, 1388–1397 (2013).

22. C. W. Woo, L. Schmidt, A. Krishnan, M. Jepma, M. Roy, M. A. Lindquist, L. Y. Atlas, T. D. Wager, Quantifying cerebral contributions to pain beyond nociception. Nat Commun 8, 14211 (2017).

23. M. M. Nickel, L. Tiemann, V. D. Hohn, E. S. May, C. Gil Avila, F. Eippert, M. Ploner, Temporal-spectral signaling of sensory information and expectations in the cerebral processing of pain. Proc Natl Acad Sci U S A 119, (2022).

24. A. Strube, M. Rose, S. Fazeli, C. Buchel, The temporal and spectral characteristics of expectations and prediction errors in pain and thermoception. Elife 10, (2021).

25. L. Tiemann, E. S. May, M. Postorino, E. Schulz, M. M. Nickel, U. Bingel, M. Ploner, Differential neurophysiological correlates of bottom-up and top-down modulations of pain. Pain 156, 289–296 (2015).

26. F. P. de Lange, M. Heilbron, P. Kok, How Do Expectations Shape Perception? Trends Cogn Sci 22, 764–779 (2018).

27. K. Friston, A theory of cortical responses. Philos Trans R Soc Lond B Biol Sci 360, 815–836 (2005).

28. L. H. Arnal, A. L. Giraud, Cortical oscillations and sensory predictions. Trends Cogn Sci 16, 390–398 (2012).

29. A. M. Bastos, W. M. Usrey, R. A. Adams, G. R. Mangun, P. Fries, K. J. Friston, Canonical microcircuits for predictive coding. Neuron 76, 695–711 (2012).

30. G. Michalareas, J. Vezoli, S. van Pelt, J. M. Schoffelen, H. Kennedy, P. Fries, Alpha-Beta and Gamma Rhythms Subserve Feedback and Feedforward Influences among Human Visual Cortical Areas. Neuron 89, 384–397 (2016).

31. C. Bradley, H. Bastuji, L. Garcia-Larrea, Evidence-based source modeling of nociceptive cortical responses: A direct comparison of scalp and intracranial activity in humans. Hum Brain Mapp 38, 6083–6095 (2017).

32. M. Vinck, R. Oostenveld, M. van Wingerden, F. Battaglia, C. M. Pennartz, An improved index of phase-synchronization for electrophysiological data in the presence of volume-conduction, noise and sample-size bias. Neuroimage 55, 1548–1565 (2011).

33. A. Mouraux, J. M. Guerit, L. Plaghki, Non-phase locked electroencephalogram (EEG) responses to CO2 laser skin stimulations may reflect central interactions between A-delta- and C-fibre afferent volleys. Clin Neurophysiol 114, 710–722 (2003).

34. J. Gross, A. Schnitzler, L. Timmermann, M. Ploner, Gamma oscillations in human primary somatosensory cortex reflect pain perception. PLoS Biol 5, e133 (2007).

35. L. Hu, W. Peng, E. Valentini, Z. Zhang, Y. Hu, Functional features of nociceptive-induced suppression of alpha band electroencephalographic oscillations. J Pain 14, 89–99 (2013).

36. Z. G. Zhang, L. Hu, Y. S. Hung, A. Mouraux, G. D. Iannetti, Gamma-band oscillations in the primary somatosensory cortex - a direct and obligatory correlate of subjective pain intensity. J Neurosci 32, 7429–7438 (2012).

37. P. Fries, Rhythms for Cognition: Communication through Coherence. Neuron 88, 220–235 (2015).

38. L. A. Baccala, K. Sameshima, Partial directed coherence: a new concept in neural structure determination. Biol Cybern 84, 463–474 (2001).

39. S. Geuter, S. Boll, F. Eippert, C. Buchel, Functional dissociation of stimulus intensity encoding and predictive coding of pain in the insula. Elife 6, (2017).

40. T. Egner, J. M. Monti, C. Summerfield, Expectation and surprise determine neural population responses in the ventral visual stream. J Neurosci 30, 16601–16608 (2010).

41. C. Keysers, V. Gazzola, E.-J. Wagenmakers, Using Bayes factor hypothesis testing in neuroscience to establish evidence of absence. Nature Neuroscience 23, 788–799 (2020).

42. M. Allen, D. Poggiali, K. Whitaker, T. R. Marshall, R. A. Kievit, Raincloud plots: a multi-platform tool for robust data visualization. Wellcome Open Res 4, 63 (2019).

43. M. Ploner, J. Gross, L. Timmermann, B. Pollok, A. Schnitzler, Pain suppresses spontaneous brain rhythms. Cereb Cortex 16, 537–540 (2006).

44. L. Tiemann, V. D. Hohn, S. Ta Dinh, E. S. May, M. M. Nickel, J. Gross, M. Ploner, Distinct patterns of brain activity mediate perceptual and motor and autonomic responses to noxious stimuli. Nat Commun 9, 4487 (2018).

45. S. Albu, M. W. Meagher, Expectation of nocebo hyperalgesia affects EEG alpha-activity. Int J Psychophysiol 109, 147–152 (2016).

46. N. T. Huneke, C. A. Brown, E. Burford, A. Watson, N. J. Trujillo-Barreto, W. El-Deredy, A. K. Jones, Experimental placebo analgesia changes resting-state alpha oscillations. PLoS One 8, e78278 (2013).

47. C. W. Woo, M. Roy, J. T. Buhle, T. D. Wager, Distinct brain systems mediate the effects of nociceptive input and self-regulation on pain. PLoS Biol 13, e1002036 (2015).

48. M. Zunhammer, U. Bingel, T. D. Wager, C. Placebo Imaging, Placebo Effects on the Neurologic Pain Signature: A Meta-analysis of Individual Participant Functional Magnetic Resonance Imaging Data. JAMA Neurol 75, 1321–1330 (2018).

49. M. N. Baliki, B. Petre, S. Torbey, K. M. Herrmann, L. Huang, T. J. Schnitzer, H. L. Fields, A. V. Apkarian, Corticostriatal functional connectivity predicts transition to chronic back pain. Nat Neurosci 15, 1117–1119 (2012).

50. A. Mansour, A. T. Baria, P. Tetreault, E. Vachon-Presseau, P. C. Chang, L. Huang, A. V. Apkarian, M. N. Baliki, Global disruption of degree rank order: a hallmark of chronic pain. Sci Rep 6, 34853 (2016).

51. S. Ta Dinh, M. M. Nickel, L. Tiemann, E. S. May, H. Heitmann, V. D. Hohn, … M. Ploner, Brain dysfunction in chronic pain patients assessed by resting-state electroencephalography. Pain, (2019).

52. H. Heitmann, C. Gil Avila, M. M. Nickel, S. Ta Dinh, E. S. May, L. Tiemann, V. D. Hohn, T. R. Tolle, M. Ploner, Longitudinal resting-state electroencephalography in patients with chronic pain undergoing interdisciplinary multimodal pain therapy. Pain, (2022).

53. Y. Zhang, W. Wu, R. T. Toll, S. Naparstek, A. Maron-Katz, M. Watts, … A. Etkin, Identification of psychiatric disorder subtypes from functional connectivity patterns in resting-state electroencephalography. Nat Biomed Eng, 309–323 (2021).

54. R. Kuner, H. Flor, Structural plasticity and reorganisation in chronic pain. Nat Rev Neurosci 18, 113 (2017).

55. C. Pernet, M. I. Garrido, A. Gramfort, N. Maurits, C. M. Michel, E. Pang, … A. Puce, Issues and recommendations from the OHBM COBIDAS MEEG committee for reproducible EEG and MEG research. Nat Neurosci 23, 1473–1483 (2020).

56. F. Faul, E. Erdfelder, A. G. Lang, A. Buchner, G*Power 3: a flexible statistical power analysis program for the social, behavioral, and biomedical sciences. Behav Res Methods 39, 175–191 (2007).

57. J. Correll, C. Mellinger, G. H. McClelland, C. M. Judd, Avoid Cohen’s ‘Small’, ‘Medium’, and ‘Large’ for Power Analysis. Trends Cogn Sci 24, 200–207 (2020).

58. A. S. Zigmond, R. P. Snaith, The hospital anxiety and depression scale. Acta Psychiatr Scand 67, 361–370 (1983).

59. I. Bjelland, A. A. Dahl, T. T. Haug, D. Neckelmann, The validity of the Hospital Anxiety and Depression Scale. An updated literature review. J Psychosom Res 52, 69–77 (2002).

60. L. Hu, G. D. Iannetti, Neural indicators of perceptual variability of pain across species. Proc Natl Acad Sci U S A, (2019).

61. T. P. Jung, S. Makeig, C. Humphries, T. W. Lee, M. J. McKeown, V. Iragui, T. J. Sejnowski, Removing electroencephalographic artifacts by blind source separation. Psychophysiology 37, 163–178 (2000).

62. R. Oostenveld, P. Fries, E. Maris, J. M. Schoffelen, FieldTrip: Open source software for advanced analysis of MEG, EEG, and invasive electrophysiological data. Comput Intell Neurosci 2011, 156869 (2011).

63. B. D. Van Veen, W. van Drongelen, M. Yuchtman, A. Suzuki, Localization of brain electrical activity via linearly constrained minimum variance spatial filtering. IEEE Trans Biomed Eng 44, 867–880 (1997).

64. D. J. Thomson, Spectrum Estimation and Harmonic Analysis. Proceedings of the IEEE 70, 1055–1096 (1982).

65. J. N. Rouder, R. D. Morey, Default Bayes Factors for Model Selection in Regression. Multivariate Behav Res 47, 877–903 (2012).

66. J. van Doorn, A. Ly, M. Marsman, E. J. Wagenmakers, Bayesian rank-based hypothesis testing for the rank sum test, the signed rank test, and Spearman’s rho. Journal of Applied Statistics 47, 2984–3006 (2020).

67. C. M. Bishop, Pattern recognition and machine learning. Information science and statistics (Springer, New York, 2006), pp. xx, 738 p.

